# Tunable DNMT1 degradation reveals cooperation of DNMT1 and DNMT3B in regulating DNA methylation dynamics and genome organization

**DOI:** 10.1101/2023.05.04.539406

**Authors:** Andrea Scelfo, Viviana Barra, Nezar Abdennur, George Spracklin, Florence Busato, Catalina Salinas-Luypaert, Elena Bonaiti, Guillaume Velasco, Anna Chipont, Coralie Guérin, Andréa E. Tijhuis, Diana C.J. Spierings, Claire Francastel, Floris Foijer, Jorg Tost, Leonid Mirny, D. Fachinetti

## Abstract

DNA methylation (DNAme) is a key epigenetic mark that regulates critical biological processes maintaining overall genome stability. Given its pleiotropic function, studies of DNAme dynamics are crucial, but currently available tools to interfere with DNAme have limitations and major cytotoxic side effects. Here, we present untransformed and cancer cell models that allow inducible and reversible global modulation of DNAme through DNMT1 depletion. By dynamically assessing the effects of induced passive demethylation through cell divisions at both the whole genome and locus-specific level, we reveal a cooperative activity between DNMT1 and DNMT3B to maintain and control DNAme. Moreover, we show that gradual loss of DNAme is accompanied by progressive and reversible changes in heterochromatin abundance, compartmentalization, and peripheral localization. DNA methylation loss coincided with a gradual reduction of cell fitness due to G1 arrest, but with minor level of mitotic failure. Altogether, this powerful system allows DNMT and DNA methylation studies with fine temporal resolution, which may help to reveal the etiologic link between DNA methylation dysfunction and human disease.

## INTRODUCTION

Epigenetic modifications are master regulators of the chromatin dynamics. They influence the association to DNA and downstream functions of binding factors, ultimately leading to tight control of various biological processes such as gene expression and chromatin conformation. The fundamental role of epigenetic information is highlighted by the fact that it is heritable across cell divisions.

DNA methylation (DNAme) was the first epigenetic modification described in mammals [1]. It is accomplished by the covalent binding of a methyl moiety transferred from S-adenosyl-L-methionine (SAM) to the fifth carbon of cytosine (C) in the context of CpG dinucleotides. This enzymatic reaction generates the 5-methylcytosine (5mC). 5mC, which accounts for 4% of all cytosines [2], is frequently referred to as “the fifth base”; typically, 80% of CpGs in mammalian genomes are methylated. Nevertheless, about 60% of mammalian genes have promoters with high CpG density, called CpG islands (CGIs), which are normally devoid of methylation [3]. Methylated promoter CGIs also exist: it is the case of long-term repressed genes, such as germline and imprinted genes [4]. Somatic DNAme domains are erased right after fertilization to establish a totipotent germiline epigenotype, deposited *de novo* during early development and undergo massive re-shaping during differentiation, lineage specification, and in response to external cues; then, they are maintained and inherited through cell divisions [5]. DNAme is also an effective way to maintain genome stability by keeping silent the expression and transposition activity of DNA repetitive elements [6]. *De novo* deposition of methyl groups is catalyzed by DNA methyltransferases (DNMT) 3A, 3B (and 3C in mouse), and maintained by DNMT1 [7]. However, these distinct enzymatic functions are not strictly divided between DNMTs as DNMT3A and 3B may also be necessary for preservation of already-established methylation patterns [8, 9], and DNMT1 can methylate *de novo* in certain conditions, like at transposable elements, and in cancer [10–13]. However, whether this is accomplished directly by DNMT1 or by interacting partners is still debated [14].

Like other epigenetic modifications, DNAme is a reversible process. Two main de-methylating pathways have been proposed: active and passive. The active DNA demethylation pathway involves the step-wise enzymatic conversion of 5mC to 5hmC mediated by TET (Ten Eleven Translocation) enzymes, which iteratively oxidise 5mC to 5-carboxylcytosine (5caC) [15]. 5caC is then the substrate for thymidine DNA glycosylase (TDG)-mediated base excision repair, ultimately leading to unmodified C [16]. Another proposed active demethylation process relies on the deamination activity of 5mC by the AID/APOBEC family members [17]. In contrast, passive loss of DNAme does not rely on any enzymatic activity; rather, it is caused by dilution of 5mC through rounds of DNA replication in the absence of a maintenance activity.

DNAme is essential for vertebrate physiology as it regulates, directly or indirectly, different genome functions, such as gene expression, silencing of transposons, chromatin condensation, imprinting and the inactivation of the X-chromosome, overall maintenance of genome stability [18], and correct development [19]. Severe pathologies and genetic syndromes are associated with altered functions of DNMTs [20]; for example, mutations in DNMT3B cause ICF (Immunodeficiency, Centromeric instability and Facial anomalies syndrome), a rare developmental disease [21] and alterations of imprinted DNAme patterns during development cause Prader–Willy and Angelman syndromes [22], other two genetic diseases. Furthermore, aberrant DNA methylation domains were the first epigenetic anomaly observed in human cancer [23], where hypermethylation of the promoters of specific tumor suppressor genes is associated with transcriptional silencing, while repetitive sequences are massively hypomethylated [24].

The major function of DNAme is to regulate transcription by acting on gene promoters and gene bodies and/or by influencing the formation and maintenance of heterochromatin domains, which are mainly marked by H3K9me3 [25]. In addition, the presence of histone tail modifications can shape genomic DNAme patterns by controlling the recruitment of DNMTs to the DNA [26]. DNAme can contribute to the establishment of chromatin states by influencing the binding of transcriptional regulators to gene promoters and the function of chromatin remodelers [27]. Furthermore, DNAme may play a key role in genome organization as perturbing DNMTs drastically remodels chromosome compartmentalization [28], and DNAme enhances chromatin compaction and stiffness in *in vitro* studies [29]. CpGme also impacts nucleosome positioning and the mechanical and physical properties of DNA [30]. However, due to the complexity of genome organization and histone modification patterns, how DNAme affects chromatin organization over time is still unknown.

DNMT1 is necessary for long term cell survival and its depletion results in severe mitotic defects [31]. However, DNMT knock-down/out experiments have resulted in either inefficient protein depletion, or the generation of smaller isoforms retaining methylation activity [32–34]. Consequently, such systems have been unable to fully elucidate the roles of DNMT1 and DNAme maintenance and lack temporal detail. As an alternative, demethylating agents (DNMTs inhibitors such as 5-azacytidine (AZA) and 2’deoxy-5-azacytidine (DAC)) have been used to investigate DNAme, but these cause unavoidable secondary cytotoxic effects [35]. Alternative strategies achieve targeted demethylation using a fusion of the TET enzyme and inactive Cas9 (dCas9) [36, 37] or the steric hindrance created by dCas9 on the DNA preventing DNMTs binding [38]. However, these approaches only achieve locus-specific demethylation. An inducible and efficient system that drives global DNA demethylation with limited side effects has not yet been developed.

Here, to study DNAme dynamics and its impact on cellular physiology, we generated cellular models for tunable DNAme in which the DNMT1 enzyme can be inducibly degraded in a rapid and efficient manner through a degron system approach. Using this system, we analyze the dynamics and consequences on chromatin organization and cell fitness of passive DNA demethylation through cell divisions without the toxicity caused by demethylating drugs. To further study the interdependence between DNA methyltransferase activities, we also engineered the DNMT1 degron system in a DNMT3B^-/-^ genetic background. Here, we evaluate time-dependent changes in methylation patterns at a genome-wide level upon DNMT1 depletion and the effect on cell proliferation and chromatin organization. We also show that DNMT1 and DNMT3B can cooperate to establish and maintain CpG methylation and chromatin domain interactions in a somatic context. The advantages of this system—rapid degradation of DNMT1 leading to massive DNA demethylation, reversibility, and low impact on cell cycle progression—make these cell lines suitable for accurate DNA methylation studies.

## RESULTS AND DISCUSSION

### Generation of an inducible DNMT1 degradation system

To study DNAme dynamics, we generated a pseudo-diploid colorectal carcinoma epithelial (DLD-1) and a non-transformed, diploid retinal pigment epithelial cell line (RPE-1) in which the endogenous DNMT1 protein can be rapidly degraded through the auxin (indol-3-acetic acid, IAA)-inducible degron system (AID) (Figure 1A) [39, 40]. To engineer a miniAID (A) module fused with the fluorescent protein mNeonGreen (N) at the endogenous *DNMT1* locus, we took advantage of CRISPR-Cas9 genome editing (Figure S1A). Single cell clones were isolated, and the edited locus was verified by PCR (Figure S1B) and sequencing. Cell imaging analysis and immuno-blot (Figure 1B,E and S1C, D) showed efficient DNMT1 degradation in both homozygous (*^NA/NA^DNMT1*) and heterozygous clones (*^+/NA^DNMT1*) within 2 hours of IAA addition (Figure 1F, G).

**Figure 1.**
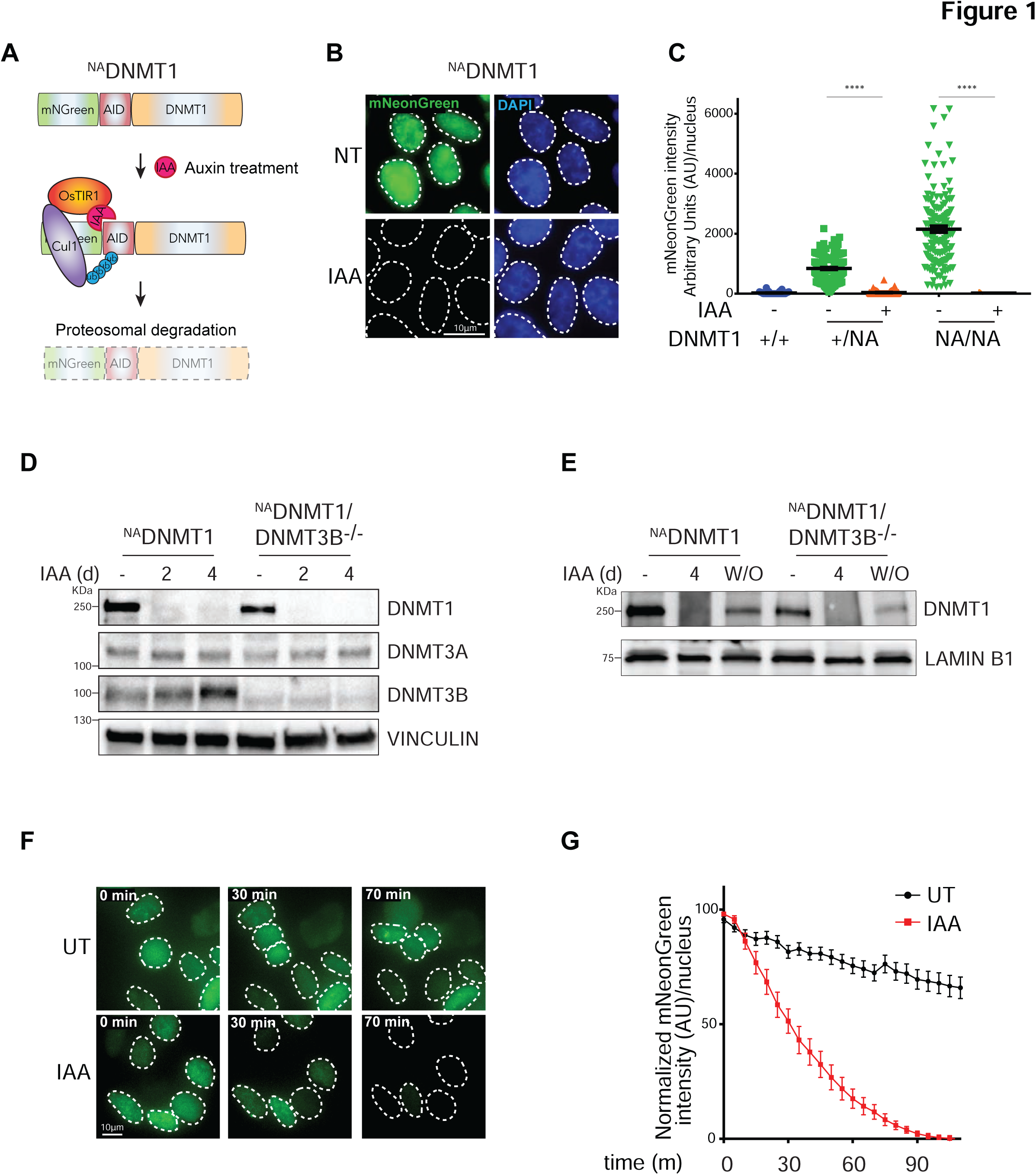
Inducible, rapid and complete DNMT1 degradation. **A.** Schematics of the endogenous DNMT1 gene tagging strategy to achieve protein degradation. AID: Auxin Inducible Degron; OsTIR1: *Oryza Sativa* TIR1; Cul1: Ubiquitin Ligase; Ub: ubiquitin. IAA: indol-3-acetic acid. **B.** Representative immunofluorescence analysis showing mNeonGreen-tagged DNMT1 signal (NT) and its degradation upon IAA treatment (24 h). Scale bar, 10µm. **C.** Quantification of nuclear mNeonGreen signal (DNMT1) in the indicated DLD-1 cell clones in untreated and IAA-treated condition (24h). +/+: wild type; +/NA: heterozygous *DNMT1*-tagged clone; NA/NA: homozygous clone. Error bar represents the SEM; each dot represents one analyzed nucleus (N > 100 per conditions). Unpaired t test: ****p <0.0001. **D.** DNMTs protein levels in the indicated cell lines treated with IAA for 2 and 4 days compared to an untreated control. VINCULIN served as loading control. **E.** Immunoblot analysis showing DNMT1 re-accumulation after IAA washout (W/O, 4 days) in the indicated cell lines. LAMINB1 served as loading control. **F-G.** Representative live cell imaging (F) and relative quantification analysis (G) showing rapid DNMT1 depletion at the indicated times from IAA treatment. Error bar represents the SEM. Scale bar, 10µm.

Upon IAA-induced DNMT1 degradation in a homozygous clone (hereinafter named ^NA^DNMT1) for 4 days, we observed upregulation of DNMT3B, while no changes were observed in the overall DNMT3A levels (Figure 1D). To counteract this phenomenon and properly model the dynamics of DNA demethylation, we knocked-out *DNMT3B* in the ^NA^DNMT1 DLD-1 background. Correct genetic DNMT3B deletion and DNMT1 degradation was confirmed by WB analysis and imaging (Figure 1D, S1E).

We then took advantage on another key feature of the AID system, its reversibility: after auxin wash-out (W/O) we can indeed observe re-accumulation of DNMT1 within 2-4 days, although to a lesser extent in the *^NA^DNMT1/DNMT3B^-/-^*cell line (Figure 1E).

### DNMT1 and DNMT3B cooperate to control DNA methylation dynamics at different genomic loci

As DNMT1 is the major enzyme involved in DNA methylation maintenance, its IAA-dependent degradation is expected to result in a progressive loss of DNAme across cycles of DNA replication.

To evaluate the degree of global DNA demethylation upon DNMT1 degradation across cell generations, we performed DNA-based assays to analyse the 5-methyl-Cytosine (5mC) levels at different time points after IAA addition (Figure 2A). Global DNAme scored by dot-blot analysis showed decreased levels in *^NA^DNMT1* and *^NA^DNMT1/DNMT3B^-/-^* cell lines upon IAA administration over time (Figure S2A). We confirmed these results using immuno-fluorescence (Figure S2B, C). Of note, DNMT1 conditional depletion led to a greater reduction in global DNAme than treatment with the demethylating agent (5-aza-2’-deoxycytidine, DAC) (Figure S2A, S2D), a widely used demethylating agent that leads to DNA damage and cytotoxicity [41].

**Figure 2.**
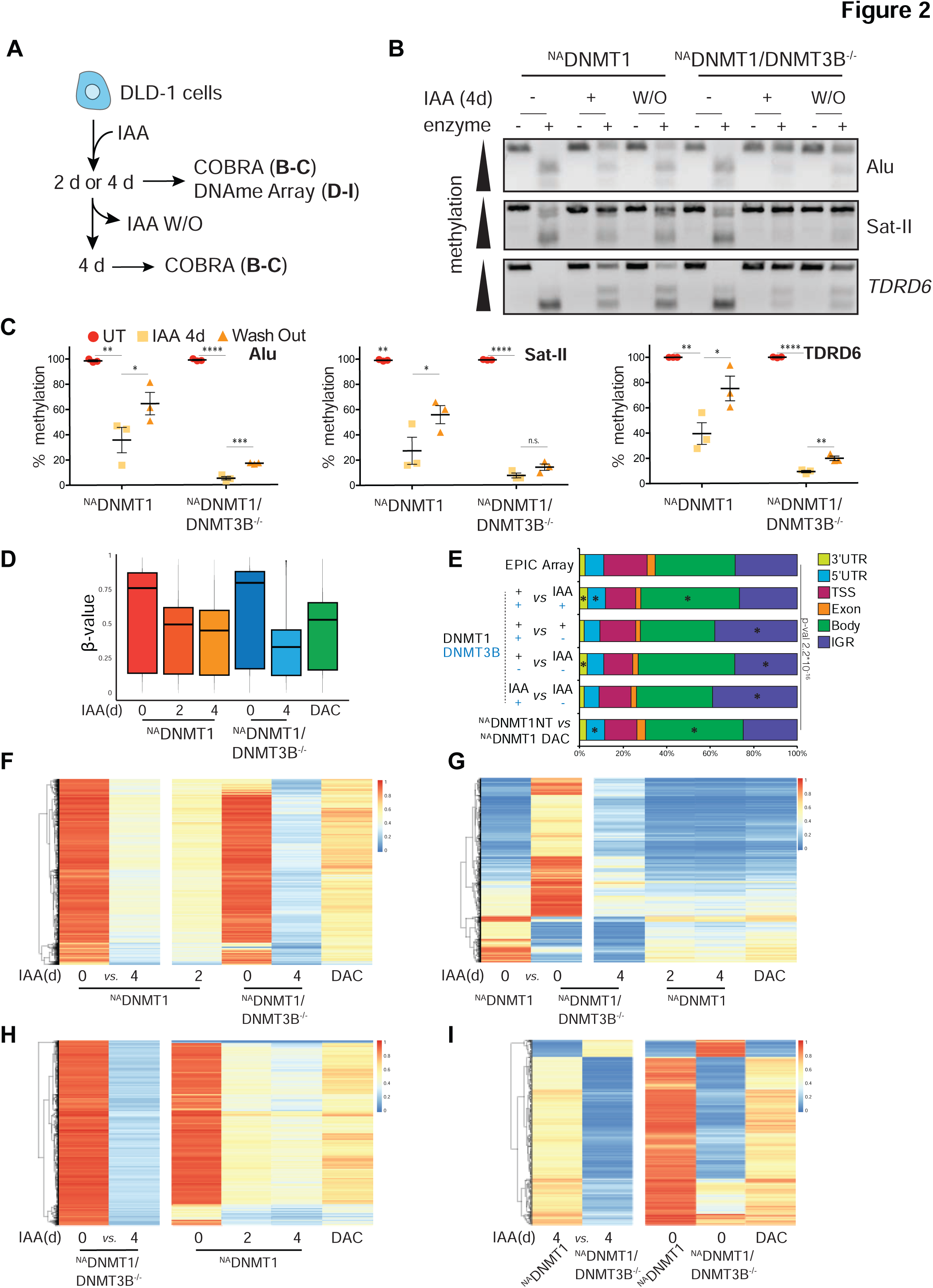
Induced DNMT1 degradation leads to progressive DNA demethylation. **A**. Schematics of the experiments shown in panels B-I. **B.** Representative agarose gel of Combined Bisulfite Restriction Analysis (COBRA) at selected loci in the indicated cell lines and treatments. BstUI and BstBI enzymes were used for *TDRD6*, Alu and SatII, respectively. W/O = washout. **C.** Quantification of methylated DNA (percentage of the total) of COBRA assay normalized to relative untreated control. Each dot represents one biological replicate (N=3); error bars represent the SEM. Unpaired t test: Alu:*p=0.049, **p= 0017, ***p=0.006; Sat-II: *p=0.0456, **p= 0013; *TDRD6:* *p=0.0255, **p= 0.001 and 0.003; ****p <0.0001. **D.** Boxplot of methylation β-values of probes identified by the EPIC Array in the indicated cell line and conditions. Dark line indicated the medians; boxes indicate Q1 and Q3; whiskers extend to include 99% of the data. **E.** Genome annotation analysis of the differentially-methylated probes (DMPs) identified in the indicated pairwise comparisons. The distribution of the EPIC Array probes is shown as reference. UTR: untranslated region; TSS: Transcription Start Site IGR: inter-genic regions. Chi-square test was used to calculate p-value and define significant changes in the distribution of DMPs of the indicated categories relative to EPIC array composition. Stars (*) indicate the genomic loci with major change (i.e. contributing the most to the p-value based on the standardized residuals, stdres> 2). **F-I.** Heatmap showing methylation intensity of DMPs (p <0.05, Δβ-value≥30%) among the indicated cell lines and conditions. The first two columns represent the DMPs identified in the indicated pairwise comparison. The methylation status of the same DMPs in the other cell lines and treatment is also shown. Due to the limited capacity of graphical visualization of high number of DMPs, in (F) are shown 40,697 DMPs (Δβ-value≥40%) out of 106,647 (Δβ-value≥30%) as indicated in the text; in (G) are shown 18,874 DMPs DMPs (Δβ-value≥40%) out of 46739 (Δβ-value≥30%) as indicated in the text; similarly, in (H) are shown 12,381 DMPs (Δβ-value≥60%) out of 455,309 as stated in the text (Δβ-value≥30%).

We next scored the degree of DNMT1 degradation-induced demethylation at selected genomic loci by COBRA (Combined Bisulfite Restriction Analysis) [42] and MeDIP (Methylated DNA Immunoprecipitation) analyses [43]. With these assays, we observed locus-specific methylation reduction at 48 and 96 hours after IAA treatment, with some regions being more susceptible than others (Figure 2B, C). COBRA analysis showed significant demethylation at Alu and Satellite II (Sat II, chromosome 1; accession number X72623) repetitive sequences and at the Tudor Domain Containing 6 (*TDRD6*) promoter, a germline gene carrying a high DNA methylation load in differentiated cells [44]. Four days after IAA withdrawal (W/O), loci partially regain their methylation, but mainly in the *DNMT3B* wild-type background (Figure 2B, C). Analogous results were obtained by MeDIP analysis of the Farnesoid X (*FXR*) and the Collagen allpha2 (*COL1A2*) transcription start sites, which bear different degrees of CpG island methylation in DLD-1 cells (Figure S2E). In this case, DNAme was fully recovered after IAA withdrawal even in *^NA^DNMT1/DNMT3B^-/-^* cells, possibly because of partial DNAme reduction of these loci.

To broaden our results on DNAme loss, we performed a genome-wide analysis through an Infinium Methylation EPIC array. This approach allows us to investigate DNAme quantitatively across over 850000 CpG sites. We compared DNMT1 depletion at 2 and 4 days in a wild-type and DNMT3B^-/-^ genetic background compared to a non-treated control and DAC treatment in DLD-1 cells. Principal Component Analysis (PCA) shows different distribution among conditions highlighting altered global DNA methylation pattern of DNMT3B^-/-^ cells compared to wild type, with the major changes observed in the *^NA^DNMT1/DNMT3B^-/-^*after 4 days of depletion. DAC-treated cells for 4 days are similar to *^NA^DNMT1* cells after 2 days IAA-mediated DNMT1 depletion for 2 days (Figure S2F). Probes’ methylation value (β-value) shows that DNAme is gradually lost with time upon IAA-mediated DNMT1 degradation at a higher level compared to DAC treatment (Figure 2D). Surprisingly, the absence of DNMT3B slightly increases bulk DNAme level, which is reduced upon 4 days of DNMT1 depletion (Figure 2D).

To investigate the role and inter-dependence of DNMT1 and DNM3B in regulating DNAme, we identified the differentially methylated probes (DMPs) between two conditions (adjusted pval<0.05 and Δβ>0.3). By comparing the analyzed DMPs to probes’ distribution on the EPIC array, DNMT1 depletion causes DNAme loss mostly in gene bodies, while DNAme loss at intergenic regions are mostly susceptible to the absence of DNMT3B (Figure 2E). Upon 2 and 4 days of DNMT1 depletion via IAA addition, 106,647 (“early”) and 178,529 probes, respectively, show at least 30% of methylation loss with respect to the untreated control (Figure 2F, S2G). This timing corresponds with the time required to achieve passive demethylation via the DNA replication cycle. Of these 178,529 probes, 79,617 lose methylation only after 4 days of IAA treatment (“late”). The total demethylation achieved using this system is approximately 4 times higher than that obtained with DAC treatment (43,790 hypomethylated probes with Δβ>0.3 compared to untreated control), highlighting the advantage of our genetic system to study DNAme dynamics over classical demethylating agents. Importantly, most of these DMPs did not lose DNAme in a *DNMT3B* null context, but the degree of demethylation at these sites is stronger in this KO background upon 4 days of DNMT1 depletion, suggesting a cooperative effect of DNMT1 and DNMT3B in maintaining DNAme (Figure 2F).

To examine the role of DNMT3B in maintaining DNAme, we compared the untreated *^NA^DNMT1* with *^NA^DNMT1/3B^-/-^* cell lines (Figure 2G). We identified a total of 46,739 DMPs (adj. pval<0.05), and only 30% of them (13,958) lose methylation in the *DNMT3B* KO vs. wildtype. Indeed, 70% of the identified DMPs (32,781) display an increased DNAme status compared to control; this increased methylation is lost upon 4 days of DNMT1 depletion (Figure 2G). This DNMT1-dependent DNAme suggests an uncontrolled DNMT1 activity in the absence of DNMT3B, which is not simply due to DNMT1 upregulation in *^NA^DNMT1/3B^-/-^* cells (Figure 1D).

Next, we assessed the contribution of DNMT1 to DNAme in the absence of DNMT3B (Figure 2H). DNMT1 depletion in the *DNMT3B^-/-^* genetic background has the greatest effect on demethylation, with 455,309 total DMPs (pval<0.05 and Δβ>0.3). These probes bear higher methylation levels when DNMT3B is expressed, indicating that DNMT1 and DNMT3B act cooperatively. Even at these DMPs, DAC is less efficient in inducing CpGs demethylation. With the scope to investigate the DNAme activity of DNMT3B in absence of DNMT1, we compared 4 days IAA-treated *^NA^DNMT1* vs. 4 days IAA-treated *^NA^DNMT1/3B^-/-^* cells, and we identified 5,213 DMPs (pval<0.05 and absolute Δβ 0.3) (Figure 2I). Also in this case, we observed a cooperative activity of DNMT1 and DNMT3B since 4 days IAA-treated *^NA^DNMT1/3B^-/-^* cells show higher level of DNAme loss of most of the probes (4,523) compared to untreated ^NA^DNMT1/3B^-/-^ cells (Figure 2I). Similar to what we observed in the ^NA^DNMT1/3B^-/-^ cells, among the aforementioned 5,213 DMPs we identified 690 DMPs whose methylation level is higher in the absence of DNMT3B. This could be due to residual DNMT1 activity, compensatory mechanisms, and/or to a lesser sensitivity to demethylation since their methylation load is high in untreated ^NA^DNMT1/3B^-/-^ (Figure 2I).

To investigate if the DMPs regulated by DNMT1 and DNMT3B are located in specific epigenetic contexts, we took advantage of the ChromHMM tracks that categorize the chromatin states of the human genome into fifteen categories [45]. By comparing the analyzed DMPs to probes’ distribution on the EPIC array, DNMT1 depletion preferentially induced progressive hypomethylation at facultative heterochromatin (Polycomb-repressed sites), transcription transition and elongation sites, weakly transcribed sequences and enhancer elements, while DNAme at active promoters was unaffected (Figure S2H). DNAme loss at Polycomb-repressed regions was also observed in DNMT3B^-/-^, with additional changes occurring at constitutive heterochromatin and insulators. In contrast, DNMT3B loss had no impact on transcriptionally active loci and strong enhancers and, as with DNMT1, DNAme at active promoters was unaffected (Figure S2I). In addition, using the Repeat Masker tool (www.repeatmasker.org) [46], we assessed how DNMT1 and DNMT3B affect DNAme levels at repetitive genome elements. DNMT1 has a major impact on LINE and SINE elements and DNA repeats (Figure S2J). On the other hand, while DNMT3B loss has the same effect on LINE elements, it has no impact on SINEs and DNA repeats; unlike DNMT1, we observed DNMT3B-dependent demethylation of LTRs (Figure S3K). Together, this suggests some degree of targeting of the enzymes to different types of chromatin, with DNMT3B acting more specifically in heterochromatin (according to ChromHMM and LINE-rich regions), while DNMT1 acting more broadly both euchromatin and euchromatin (as evident from ChromHMM and effect on both LINE and SINE-rich regions).

Overall, we demonstrate that the generated cell lines are a powerful tool to study DNA demethylation dynamics. We show that DNMT1 and DNMT3B, alone or simultaneously, are needed to achieve precise DNAme patterns throughout the genome. The presence of genomic loci gaining DNAme in a DNMT1-dependent manner in DNMT3B^-/-^ cells highlight a possible non canonical or uncontrolled spurious *de-novo* DNMT1 methyltransferase activity at selected genomic loci, which, based on previous *in vitro* [47, 48] and *in vivo* experiments, was proposed to be limited to transposable elements [12], non-CpG sites [49] and selected promoters [50]. As DNMT1 degradation can be reversed by IAA withdrawal, this system will be useful for examination of the re-methylation of CpGs.

### Conditional DNMT1 depletion partially impairs cell proliferation, while co-depletion with DNMT3B leads to cell lethality

To assess the effect of gradual DNA demethylation on cell proliferation and survival without the secondary effects of chemical demethylating agents [35], we performed cell cycle analysis in p53-deficient DLD-1 *^NA^DNMT1* and *^NA^DNMT1/3B^-/-^* cells with up to 10 days of IAA-mediated DNMT1 depletion (Figure 3A, S3A). DNMT1 depletion over time led to an impaired cycling rate; after 8-10 days of IAA treatment, we observed an increased percentage of cells in G1 and reduced S-phase (Figure 3B). These defects in the cell cycle progression were further enhanced in the absence of DNMT3B: here, cells start to accumulate in G1 after just 4 days of depletion, and the impact on S phase is more pronounced. DNMT1/3B co-depleted cells, but not single DNMT1 depleted cells, undergo to cell death, suggested by an increased percentage of sub-G1 positive cells (Figure 3A, S3A). Following DNMT1 depletion, the arrest in G1 occurred faster in p53-proficient diploid RPE-1 cells compared to oncogene-positive DLD-1, suggesting a rapid activation of the cell cycle checkpoint (Figure S3B).

**Figure 3.**
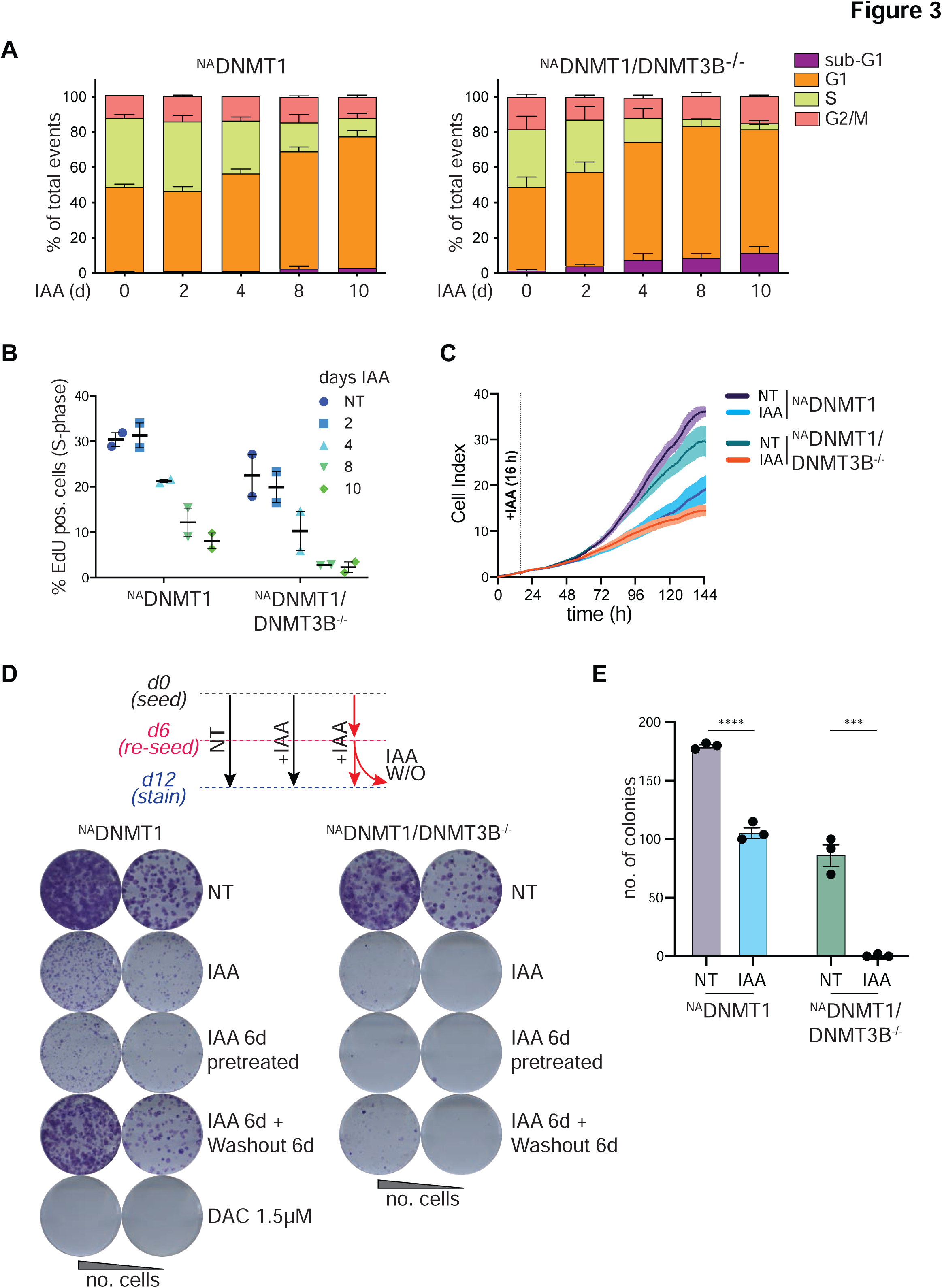
DNMTs degradation impairs cell proliferation. **A**. Cell cycle profiling by EdU/DAPI incorporation upon induced DNMT1 degradation at the indicated days from IAA treatment in *^NA^DNMT1* (left) and *^NA^DNMT1/DNMT3B^-/-^*cells. N=2, Error bars represent SEM. **B.** Quantification of S-phase (% over the total) from A in the indicated cell lines and treatments. Dots represent biological replicates; bars represent mean with SEM. **C.** Real-time Cell Index measurement of the indicated cell lines upon IAA treatment. Measurements were normalized to the 16h time point, time of IAA addition after seeding. Curves represent the mean Cell Index value from triplicates ± SD. **D-E.** Representative images of colony formation assay and relative schematics and quantifications performed in the indicated cell lines and treatments. Number of obtained colonies in selected conditions are plotted in E. N=3, Error bars represent SEM. Unpaired t test: ****p <0.0001; ***p=0.002

To assess the consequences of DNMT1 and/or DNMT3B depletion on cell proliferation, we performed no dye, live cell imaging analysis for 6 days on IAA-treated or untreated cells. While DNMT1 or DNMT3B single-depleted cells show a lower proliferation rate than wild type cells, simultaneous depletion of DNMT1 and DNMT3B has a major effect on cell fitness (Figure 3C). The long-term effects of DNMT1 depletion in wild type and *DNMT3^-/-^* cells were then tested by colony formation assay after 12 days of IAA treatment. While 12 days of DNMT1 depletion in *^NA^DNMT1* cells has a mild but detectable effect on cell survival (Figure 3D, E), 6 days of IAA pre-treatment before seeding drastically impairs colony formation and survival. In agreement with the observed faster G1 arrest, the inhibition of cell proliferation following DNMT1 depletion is stronger in the *^NA^DNMT1* RPE-1 cell line (Figure S3C), caused by proficient p53 activity. Treatment with the hypomethylating agent DAC has a strong effect on colony formation (Figure 3D) despite causing a minor reduction of DNAme respect to IAA-induced DNMT1 depletion (Figure 2D), suggesting a DNAme-independent cytotoxic effect. Recently, for chemotherapeutic purposes, a new, more tolerable, less toxic DNMT1 inhibitor was developed (GSK3685032, [51]). When tested in our system at the concentration required to induce demethylation [51], this compound still had a more profound effect on cell proliferation than did DNMT1 depletion by IAA treatment alone (Figure S3D), suggesting some level of toxicity DNAme-independent. In agreement with the live cell imaging results, DNMT1 and DNMT3B co-depletion halts cell proliferation. In addition, the reduced proliferation potential observed after single DNMT1 depletion is rescued by IAA removal, but do not recover in the DNMT3B KO background (Figure 3D, E). However, high numbers of mitotic errors are unlikely to be the cause of the reduced proliferation potential observed in *^NA^DNMT1/3B^-/-^* IAA-treated cells, as copy number variation analysis by single cell sequencing reveals very low levels of aneuploidy compared to untreated *^NA^DNMT1/3B^-/-^* cells (20% vs 12.2%; Figure S3E).

Taken together, these data demonstrate that DNMT1 is necessary to maintain proper cell proliferation. In sharp contrast to previous reports using a inducible *DNMT1* KO in HCT116 colorectal cancer cells [31], we find that complete DNMT1 depletion does not lead to rapid and severe cell lethality and/or mitotic failure. We hypothesize that the severe cell cycle arrest and cell death seen in the previous report after only 48h of genetic deletion of selected exons of *DNMT1* gene may be due to non-specific toxic effects (e.g. from CRE recombinase expression [52]). We propose that our DNMT1 depletion system is more suitable for DNAme studies since it allows for temporal control of DNMT1 degradation. In addition, we propose that the strong additive effect on cell survival in the double DNMT1/3B depleted cells is the consequence of decreased 5mC levels as DNAme is strongly reduced in the double mutant, in agreement with DNAme being essential for cell fitness.

### DNA methyltransferase loss disrupts subnuclear compartmentalization of inactive chromatin states

DNAme is deposited at CpGs of most inactive regulatory elements that are normally confined to heterochromatin and inactive regions marked by H3K9me3 or H3K27me3, respectively [53]. Moreover, it has been proposed that DNAme can influence H3K9me3 deposition [54]. To assess the interdependence between DNAme and heterochromatin, we implemented a hybrid cytometry-microscopy (Imagestream) analysis by which we tested the subnuclear localization of such histone post-translational modifications (PTMs) upon induced demethylation (Figure 4A). H3K9me3 is typically expected to correlate with DNA density and shown to be mainly positioned at pericentromere and nuclear periphery in fully differentiated cells [55, 56]. Upon DNMT1 depletion, H3K9me3 and H3K27me3 marked regions delocalized from the nuclear periphery to the nucleoplasm (Figure 4A-C). This effect was further enhanced by simultaneous DNMT1 and DNMT3B depletion. DNMT1 re-expression rescued the localization of H3K9me3 and H3K27me3 positive loci to the nuclear periphery, but only partially in *^NA^DNMT1/3B^-/-^* cells (Figure 4A-C).

**Figure 4.**
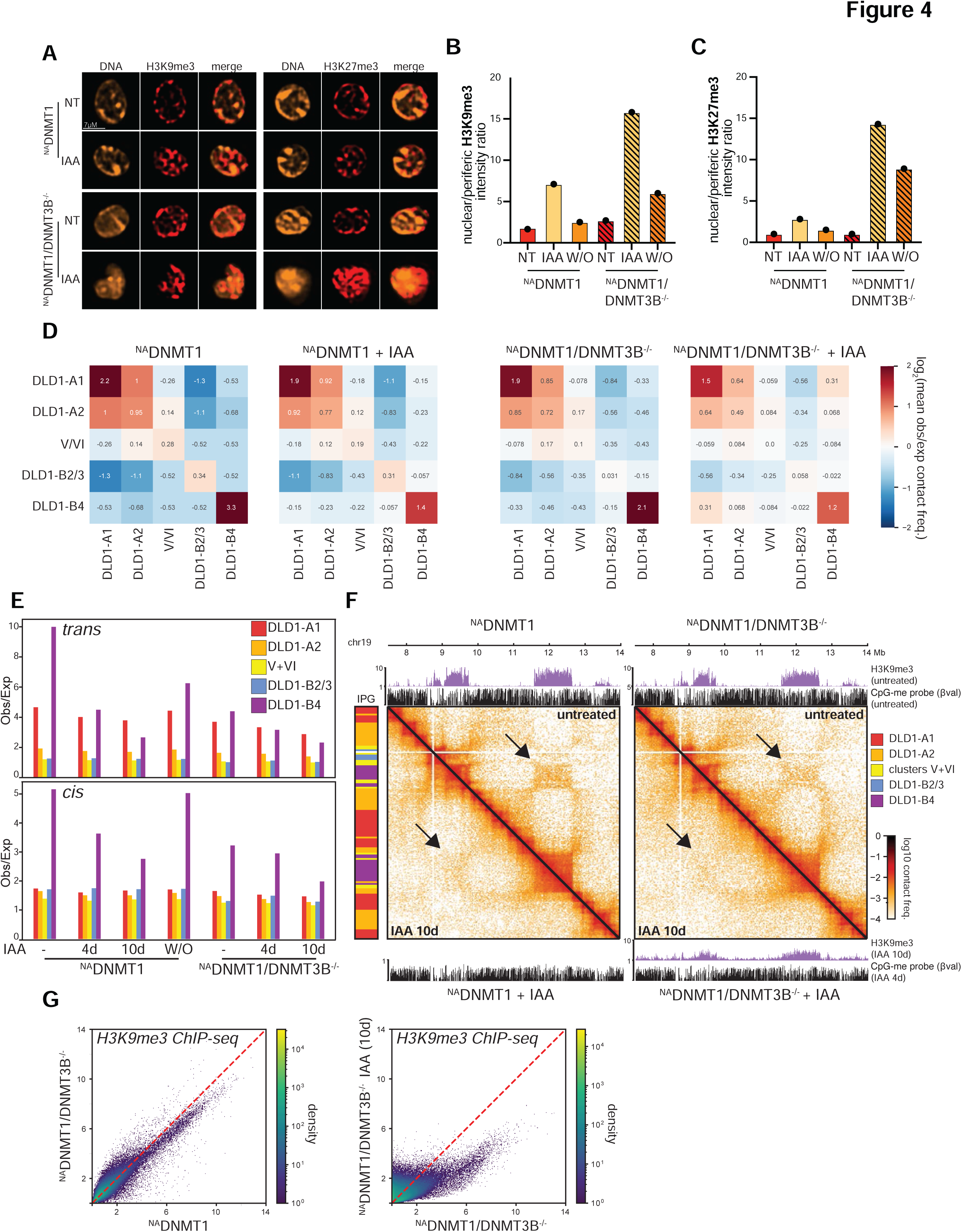
DNMTs sustain subnuclear compartmentalization of inactive chromatin. **A**. Representative immunofluorescence images of hybrid microscopy-cytometry analysis of the indicated cell lines and conditions (IAA: 10 days). Scale bar: 7 μm **B-C.** Ratio of the nuclear H3K9me3 (B) and H3K27me3 (C) signal intensities over the signal measured at the nuclear periphery by hybrid microscopy-FACS analysis in the indicated cell lines and conditions. IAA: 10 days. Washout (W/O) experiments were analyzed 4 days after IAA withdrawal (IAA: 6 days). N > 15643 cells per condition. **D.** Heatmaps of pairwise aggregate observed/expected contact frequency between interaction profile groups (IPGs), derived from 50-kb resolution contact maps for untreated and 10 days IAA-treated *^NA^DNMT1* cells and untreated and 10 days IAA-treated *^NA^DNMT1/DNMT3B^-/-^* cells, respectively. **E.** Time-courses of observed/expected contact frequency in trans (top) and in cis (bottom) within the same five defined IPGs in the indicated cell lines and conditions. The washout (W/O) experiment was analyzed 4 days after IAA withdrawal (IAA: 4 days). **F.** Example of DLD1-B4 disruption on a region of chromosome 19 in *^NA^DNMT1* (left) and ^NA^*DNMT1/DNMT3B-/-* (right) after 10 days of IAA treatment. Top to bottom: tracks of H3K9me3 (fold change over input) and CpG methylation (SWAN-normalized β−value from EPIC array) signal in untreated cells, Hi-C from untreated (top right) and treated cells (bottom left), tracks of H3K9me3 and CpG methylation from treated cells. Left: Colored track of IPG classification derived from untreated ^NA^DNMT1 cells. Arrows point to a zone of distal DLD1-B4 interaction that is visibly depleted upon treatment. **G.** Scatter plot showing loss of H3K9me3 ChIP-seq signal density (mean fold change over input) upon DNMT3B deletion (left) and DNMT1 depletion (10 days, right) in *^NA^DNMT1/DNMT3B^-/-^* cells. Each point represents a 10-kb genomic bin.

These results prompted us to investigate genome organization by chromosome conformation capture (Hi-C). We performed *in situ* Hi-C on untreated and treated (4d and 10d IAA) *^NA^DNMT1* and *^NA^DNMT1/3B^-/-^* DLD-1 cells, as well as in *^NA^DNMT1* after IAA washout. First, we compared P(s) curves, which reflect the size and density of extruded loops, and observed no significant differences between conditions, suggesting that loss of DNAme does not alter the level of cohesin-mediated loop extrusion (Figure S4A). To assess the effect of DNMT1 depletion on subnuclear compartmentalization of the genome, we next analyzed long-range contact frequencies using a recently published method [28]. While conventional compartmentalization analysis quantifies preferential interactions by imposing a binary A vs B classification, this approach partitions genomic loci based on more complex patterns of long-range interactions into groups termed subcompartments or interaction profile groups (IPGs). As in [28], our analysis disregards purely position-dependent effects on long-range interactions by consolidating clusters of loci that share a common epigenetic signature but differ in their genomic distance from the centromere (Figure S4B). Using this approach, we identified two transcriptionally active IPGs that we labeled DLD-A1 and DLD-A2 and, (following a similar naming trend as in other cell types), with the former having the strongest transcriptional activity and highest GC content. With a limited set of functional annotations, we decided not to assign a special label to clusters V and VI showing intermediate transcriptional activity. We further identified two broadly transcriptionally inactive (or heterochromatic) IPGs: DLD1-B4 and DLD1-B2/3. We assigned the label DLD1-B4 based on its epigenetic setting and positional similarity of its loci to B4 subcompartment regions in GM12878 cells [57], as well as its epigenetic similarity to the B4 IPG detected in HCT116 cells [28]. Specifically, we observe that DLD1-B4 is the only IPG enriched for the constitutive heterochromatin mark H3K9me3 and, as in GM12878 cell line, it is primarily located pericentrically and on chromosome 19 (Figure S4B).

Recently, we observed that the B4 IPG identified in HCT116 and its associated H3K9me3/HP1 chromatin state were disrupted under 5aza-mediated chemical inhibition of DNA methyltransferases and joint long-term perturbation of DNMT1 and DNMT3B using genetic knockout cells [28]. Consistent with these results, we find that DLD1-B4 compartmentalization is also disrupted under inducible DNMT1 degradation. This occurs in a time-dependent manner, reaching a roughly twofold reduction in the average observed-over-expected contact frequency between DLD1-B4 loci in both genetic backgrounds after 10 days of IAA treatment (Figure 4D-F). To validate this effect at the epigenome level, we performed ChIP-seq for H3K9me3 in untreated *^NA^DNMT1* and in untreated and 10-day treated *^NA^DNMT1/3B^-/-^*. We found that the loss of compartmentalization after treatment is accompanied by a dramatic global reduction of H3K9me3 levels (Figure 4F, G). We also observed that untreated *^NA^DNMT1/3B^-/-^* cells display weaker DLD1-B4 self-affinity and reduced H3K9me3 levels compared to untreated *^NA^DNMT1* (Figure 4D, G and S5A), suggesting that DLD1-B4 is mildly impacted by DNMT3B deletion; however, we cannot rule out if this could be due to minor differences in baseline DNMT1 levels in the two backgrounds (Figure 1D). Importantly, we observed that the impact of DNMT1 depletion on compartmentalization is reversible: after 4 days of IAA withdrawal in *^NA^DNMT1* cells treated for 4 days with IAA, DLD1-B4 self-interaction in *cis* and a recovery in *trans* recovered to ∼80% (Figure 4E, S5A).

The second inactive IPG detected in untreated *^NA^DNMT1*, which we label DLD1-B2/3, covers ∼30% of the mappable autosomal genome in these cells and has an incompletely characterized chromatin state, though it exhibits a dispersed enrichment for H3K27me3, suggesting that this state is at least partially heterochromatic (GSE85688, [58] (Figure S4B). The majority of loci of this IPG are assigned the transcriptionally inactive subcompartment labels B2 and B3 in other cell types by SNIPER (subcompartment inference using imputed probabilistic expressions) [59], a supervised learning algorithm that generalizes GM12878 subcompartment labels to other cell types (Figure S5B). Interestingly, we find that self-interaction and compartmentalization of DLD1-B2/3 are significantly disrupted in the *^NA^DNMT1/3B^-/-^* genetic background, irrespective of IAA treatment duration (Figure 4D and S5C). These results imply that a different DNA methyltransferase, DNMT3B, is required to maintain a different inactive IPG, DLD1-B2/3, suggesting a separation of function for DNMT3B and DNMT1 in genome organization.

Our Hi-C results suggest that (i) 3D compartmentalization of the DLD1-B4 IPG is progressively and reversibly sensitive to loss of DNMT1 while (ii) that of the DLD1-B2/3 IPG is sensitive to loss of DNMT3B. Additionally, depletion of DNMT1 causes the active DLD1-A1 and DLD1-A2 IPGs to exhibit decreased self-interaction preference and increased preference for interacting with inactive IPGs, indicating an overall reduction in compartmental segregation when heterochromatin is lost (Figure 4D, E). These data highlight the interplay between DNA methylation, heterochromatin, and genome organization [60, 61] and suggest that DNMTs have a significant role to play in long-range genome organization. In DLD1 cells, DNMT1 loss induces isolated but reversible changes in the preferential interaction of H3K9me3 chromatin domains found in pericentromeres and on chromosome 19. This stands in contrast to DNMT1 loss in the HCT116 cell line, where H3K9me3 domains cover a much greater proportion of the genome, thus leading to a more widespread phenotype on contact maps [28]. Conversely, we show that the preferential interaction of the chromatin state underlying the B2/3 IPG, which appears to be absent in HCT116 but is the most abundant silent IPG in DLD1 cells, depends on a different methyltransferase, DNMT3B. Taken together, our results have revealed complementary roles of different DNMTs in the maintenance of different defined chromatin states.

Our inducible system will serve as a powerful tool to elucidate in greater detail the mechanisms and temporal dynamics driving DNMT-dependent heterochromatin formation and 3D genome regulation. For example, it is notable that the DNMT1-controlled IPG domains on chromosome 19 contain clusters of Zinc finger proteins. It would be interesting to investigate if these factors are transcriptionally dependent on DNMT1 and contribute to cell homeostasis. DNMT3B depletion alone triggers loss of affinity of chromatin domains bearing features of facultative heterochromatin, probably corresponding to the chromatin delocalization from the nuclear periphery to the inner nuclear space observed by the hybrid microscopy/FACS analyses. These disrupted interactions could coincide with Lamin Associated Domains. Similarly, our system could be leveraged to study the functional link between DNAme and histone PTMs.

Finally, we note that the recovery of DLD1-B4 compartmentalization upon IAA withdrawal was rapid and near-complete (Figure S5A), while DNAme recovery is only partial (Figure 2B). This suggests that specific CpG sites may be critical, but also raises the possibility that DNMTs regulate broad heterochromatic regions in a manner that is not directly coupled to their DNA methylation activity. Similarly, other factors like proteins of the methyl-CpG-binding domain (MBD) family could contribute to heterochromatin maintenance. Dissecting these regulatory connections and how they contribute to the formation of cell-type specific heterochromatic landscapes will be the subject of future work.

## CONCLUDING REMARKS

Studies of DNAme and its dynamics have been challenging because current methods to specifically manipulate DNAme have low efficiency and high toxicity. Our newly-generated cell models enabled us to assess acute, reversible, and time-dependent effects of DNMT1 and DNAme loss on cell physiology, providing valuable information without bias due to cytotoxicity and cell adaptation to constitutive long-term protein depletion. We present rapid kinetics of DNMT1 protein depletion leading to a passive genome-wide reduction of DNAme levels. Depletion of DNMT1 can be observed already at two days after induction; however, cell lethality is far lower than with alternative drug-based systems. We show that heterochromatic regions are highly susceptible to DNMT depletion and demethylation, with loss of their localization at the nuclear periphery and disappearance of their compartmentalization patterns in Hi-C maps. Our data show that DNMT1 and DNMT3B cooperate to deposit and control DNAme patterns, and we describe a compensatory role for DNMT1 in *de novo* methylation at selected loci. Although recent efforts have been made for the generation of less toxic demethylating agents (e.g. GSK3685032, [51]), we believe that the advantages of our cellular models –reversibility, temporal control and low toxicity-, make them critical instruments for future investigation of fundamental biological questions concerning the role of DNAme and its establishment and maintenance.

## ACKNOWLEDGMENTS

We thank all the members of the Daniele Fachinetti team for discussion and Edith Pfister (UMass Chan Medical School, Worcester, MA, USA) for editing the manuscript. We thank Hélöise Muller from Ines Drinnenberg’s laboratory (Institut Curie, Paris) for advice and sharing reagents. We also acknowledge the Flow Cytometry Core Facility and the Cell and Tissue Imaging facility (PICT-IBiSA, member of the French National Research Infrastructure France-BioImaging ANR10-INBS-04) of the Institut Curie. D.F. receives salary support from the CNRS. D.F. has received support for this project by CNRS, I. Curie and “ARC labellisation program 2019”. A.S. was supported by AIRC and from the European Union’s Horizon 2020 research and innovation program under the Marie Skłodowska-Curie grant agreement No. 800924.

## AUTHOR CONTRIBUTIONS

A.S., V.B., C.S.L., E.B., obtained the experimental data; V.B. generated the cell lines; N.A., G.S. performed the Hi-C analysis under the supervision of L.M.; F.B. and J.T. performed the methylation array analysis; G.V. provided technical and analysis expertise supervised by C.F.; A.C. performed the ImageStream data analysis under the supervision of C.G.; A.E.T., D.C.J.S performed the scWGS data acquisition and analysis under the supervision of F.F.; A.S., G.S., N.A., F.B., L.M., and D.F. interpreted the results. A.S., G.S., N.A., L.M. and D.F. designed the experiments. D.F. conceptualized the project and supervised the experimental work. D.F, L.M., J.T., C.F., C.G. and F.F. obtained funding. A.S. and D.F. drafted the manuscript with input from N.A., G.S., and L.M. All authors participated in reviewing and editing the manuscript.

## MATERIAL AND METHODS

### Cell culture and treatments

Cells were cultured at 37 °C in a 5% CO_2_ atmosphere. DLD-1 and immortalized hTERT RPE-1 cells were maintained in DMEM-GlutaMAX and in DMEM:F12 media, respectively, supplemented with 10% fetal bovine serum, 100 U/mL penicillin-streptomycin, 0.13% sodium bicarbonate (RPE-1). Auxin indole-3-acetic acid sodium salt (IAA) (I5148, Sigma-Aldric) was used at 500 nM dissolved in water. 2’-deoxy-5-azacytidine (DAC) (A3656, Sigma-Aldrich) was used at 2.5 µM for 96h. GSK-3685032 (HY-139664, MedChemExpress) was used at the indicated concentrations. All cell lines were tested negative for mycoplasma contamination. IAA washout was performed by five gentle washes in PBS. Cells were then analyzed at the indicated time points.

### Plasmids and Cell line generation

The repair template containing mNeonGreen-miniAID sequence flanked by left and right homology arms (±500bp from ATG) of *DNMT1* genomic sequence was cloned into pCR-BluntII-TOPO plasmid with the Zero Blunt TOPO PCR Cloning Kit (Thermo Fisher Scientific). The *DNMT1*-targeting sgRNA (5′-GGCGGTACGCGCCGGCATCT –3′) was cloned into pX330 hSpCas9 expressing vector (Addgene #42230) and verified by sequencing.

DLD-1 and RPE-1 cells stably expressing OsTIR1-Myc9 were nucleofected with the DNMT1 targeting pX330 plasmid in presence of the repair template by electroporation using the Lonza Nucleofector with appropriate reagents (Nucleofector Solution V and L for DLD-1 and RPE-1,respectively) according to manufacturer’s instructions. A molar ratio of 9:1 between donor and the pX330 plasmid was used. Five days after transfection, mNeonGreen positive cells were FACS sorted and seeded for clone isolation. Clones were amplified and screened by PCR amplification of a region outside the homology arms (fw: 5′-GCCGCCATCGAGATGCACAG-3′; rev: 5′-CCACACACTGGGTATAGAAGTGGC-3′). Positive clones were confirmed by PCR sequencing and immunoblotting.

### Colony formation assay

Serial dilutions of cells were plated on 6-well plates and 500 nM IAA was added the day after seeding. After 14 days, colonies were fixed 10 min in methanol, washed twice with PBS 1X, stained for 10 min in in 1% crystal violet, 20% EtOH and washed twice with PBS 1X. Colonies obtained from the same serial dilution across all the analysed conditions were visually counted and plotted.

### Cell proliferation assay

Cell proliferation was measured on xCELLigence® (Agilent) E-Plate VIEW (96 wells) in an xCELLigence® eSight real-time cell analyzer. Cells were passaged and allowed to reach 70-80% confluency before being trypsinized, counted and resuspended in standard growth media to 4×10^4^ cells/mL. Blanks were measured on the E-Plates with 50 μL of standard growth media before adding 50 μL of the cell suspension, equivalent to 2000 cells, in triplicate wells per condition. Measurements were performed every 15 minutes and pictures of each well were taken every hour for a total of 6 days. After 16 hours, an additional 50 μL of media was added to all wells, either with or without IAA to a final concentration of 500 μM per well. Cell index –i.e. the electrical impedance generated from cell attachment and proliferation quantified via gold electrodes at the bottom of the cell culture plate wells-, was calculated with the RTCA eSight Software (Agilent) and normalized to the 16 h time point.

### Cell cycle analysis

EdU pulse was performed for 30 min (10 μM final), before collection and fixation in 70% Ethanol. Click reaction was carried out with homemade click chemistry buffer (Tris-HCl 100 mM, pH 8.5, CuSO4 1 mM, Azide-fluor488 5 μM, Ascorbic acid 100 mM) for 30 min at room temperature. Then, propidium iodide wad added at 2.5 μg/mL final concentration in presence of 250 μg/mL RNAseA (Thermo Fisher) and incubated at least 1h before acquisition on a LSRII Flow Cytometer (BD). Data were analyzed using FlowJo software v10 (BD). Cell doublets were excluded and the auto-gating tool was applied to select G1, S, G2/M populations.

### Single-cell DNA sequencing

One million cells were resuspended in media, washed and pelleted. Cells were resuspended in cell lysis buffer (100 mM Tris-HCl pH 7.4, 154 mM NaCl, 1 mM CaCl2, 500 µM MgCl2, 0.2% BSA, 0.1% NP-40, 10 µg/mL Hoechst 33358, 2 µg/mL propidium iodide (PI) in ultra-pure water) and incubated on ice in the dark for 15 minutes to complete lysis and generate nuclei. Resulting cell nuclei were gated for G1 phase (as determined by Hoechst and PI staining) and single nuclei were sorted into wells of 96-well plates on a MoFlo Astrios cell sorter (Beckman Coulter). 96 well plates containing nuclei and freezing buffer were stored at –80°C until further processing. Automated library preparation was then performed as previously described [62].

Libraries were sequenced on a NextSeq 500 machine (Illumina; up to 77 cycles – single end), and aligned to the human reference genome (GRCh38/hg38) using Bowtie2 (version 2.2.4 or 2.3.4.1; [63]). Duplicate reads were marked with BamUtil (version 1.0.3; [64]) or Samtools markdup (version 1.9; [65]). Resulting aligned read data were input for copy number calling using the Aneufinder R package (https://github.com/ataudt/aneufinder; [66]). GC correction was performed and artefact-prone regions (known regions of extremely high or low coverage) were blacklisted. Libraries were analysed using the dnacopy and edivisive copy number calling algorithms with variable width bins (average bin size 1 Mb, step size 500 kb). Libraries with on average less than 10 reads per bin were discarded. Additionally, results for single-cell sequencing were curated by requiring a minimum concordance of 90% between the output of the two algorithms.

### Immunoblotting

The cells were lysed with SDS-containing buffer, sonicated, BCA-quantified and denaturated at 95°C in 1X Laemmli sample buffer for 5 minutes. Samples were loaded on a 7.5% polyacrylamide gel (BioRad) and transferred to a 0,45µm nitrocellulose membrane (BioRad). The following antibodies were used for immunoblotting: DNMT1 (1:100, Santa Cruz sc-271729); DNMT3B (1:1000, Cell Signalling Technology #67259); DNM3A (1:2000, Abcam ab188470); VINCULIN (1:5000, Sigma V9264); LAMINB1 (1:1000, Proteintech 12987-1-AP).

### Immunofluorescence and live cell imaging

Cells were grown on poly-L-lysine-coated coverslips and fixed in PBS containing 4% formaldehyde and 0.1% Triton X-100 at room temperature for 10 min. Blocking was performed with 5 % Bovine Serum Albumin in 0.1 % Tween-20 in PBS for 10 min or in 2.5 % FBS (v/v), 0.2 M glycine, 0.1 % triton X-100 (v/v) in PBS for 30 min at room temperature. Primary antibodies were incubated in blocking buffer for 1 h at room temperature. After washing with PBS 0.1% Triton X-100, coverslips were incubated with secondary antibodies conjugated to fluorochromes (Jackson Immuno Research) for 45 min at RT. The following antibodies were used: DNMT3B (1:1000, Cell Signalling Technology #67259); mNeonGreen (1:500; Chromotek, 32f6). Coverslips were mounted using anti-fade reagent Prolong Gold containing DAPI (Life Technologies). Images were acquired on a DeltaVision Core system (Applied Precision) with 60X Olympus UPlanSApo oil-immersion objective (NA 1.4), 250 W Xenon light source equipped with a Photometrics CoolSNAP_HQ2 Camera. 4 μm Z-stacks were acquired (Z step size: 0.2 μm).

For 5-methyl-cytosine staining, cells were washed with 0.1% Tween-20 in PBS and fixed for 10 min in freshly prepared 4% formaldehyde in PBS. Cells were permeabilize with 0.5% Triton X-100 in PBS for 30 min and treated with 2M HCl for 30 min at 25°C. After extensive wash with 0.1% Tween-20 in PBS, cells were blocked for 1h in 2% BSA in 0.1% Tween-20 in PBS and then incubated anti-5MC antibody (1:500; A-1014 Epigentek) in 0.1% Tween-20 in PBS overnight at 4°C and subsequently after washing with 1:500 anti-mouse Cy5-conjugated in 0.1% Tween-20 in PBS for 1 h at 37°C. DNA was stained with DAPI (1 µg/mL) for 5 minutes at room temperature and cells imaged with a DeltaVision Core system (Applied Precision).

For the live-cell imaging, cells were plated on high optical quality plastic slides (Ibidi) treated 1 h before filming with siRDNA (1:1000) and imaged using a Deltavision Core system (Applied Precision).

### Image quantification

Immunofluorescence signals were quantified by using FIJI software. A mask of the nuclei was obtained by thresholding the DAPI channel and individual nuclei were detected using the Analyze Particles function. Five 15 × 15 pixel circles were drawn outside the nuclei (marked by DAPI staining) and the mean of the integrated signal was calculated (background). The integrated signal intensity of each individual nucleus was then calculated by subtracting the background.

### Hybrid microscopy-cytometry (Imagestream) analysis

Cells were collected, counted and fixed in 1% formaldehyde 20 min on ice. The reaction was blocked with final 125 mM Glycine for 3 min. After washes with 1X PBS, cells were permeabilized with 1X permeabilization buffer (PB) (kit Foxp3/Transcription Factor Staining Buffer kit, Thermo Fisher, 00-5523) for 30 min RT. Cells were resuspended in 1XPB containing H3K9me3 (1:700; Abcam, ab8898) or H3K27me3 (1:700; Cell Signalling Technology, 9733S) antibodies and incubated for 1 h at room temperature.

After three washes in 1X PB, cells were resuspended in PB with fluorochrome-conjugated secondary antibody (AlexaFluor-647, Jackson lab, 1:500) for 30 min, at room temperature. Following 2 washes in 1X PB, cells were resuspended cells in 30 µL of 1X FACS buffer (1%BSA, 2mM EDTA in PBS) + PI (10 µg/mL) and acquired on an Amnis ImageStreamX MKII device set with proper compensation and unstained controls. Lasers 405nm (70mW), 488nm (100mW), 561nm (50mW) and 642nm (mW) were used and parameters were recorded with INSPIRE software as followed: channels 01 and 09 for Brightfields (Ch01 and Ch09 BF), channel 02 for GFP (Ch02 GFP), channel 04 for Propidium iodide (Ch04 PI), channel 07 for CFP (Ch07 CFP) and channel 11 for H3K9me3 or H3K27me3 AF647 (Ch11 AF647). Data analysis was performed with IDEAS software (V6.2). Cells in focus were selected based on Gradient RMS feature on Ch01, then Singlets were isolated using Area and Aspect Ratio Intensity features on Ch01. GFP positive and Negative events were separated using Ch02 intensity, independently from CFP intensity. To focus on PI signal localization in the nucleus, three other specific masks were generated: Erode(M04, 4), consisting of eroding 4 pixels of the regular M04 mask, Dilate(M04, 1) to dilate of 1 pixel the M04 and the combined mask: Dilate(M04, 1) And Not Erode(M04, 4). The Erode(M04, 4) represents the inner part of the nucleus when the Dilate(M04, 1) and Not Erode(M04, 4) shows the nuclear membrane. Those masks were applied to an Intensity feature, allowing exclusion of outliers. To better assess the localization, an Intensity Ratio (Ratio Membrane/Intra) was then created by dividing Intensity at the membrane by the Intensity at the internal nuclear part. Nuclear (<1 “Internal”) and membrane (>1 “external”) PI were separated at Ratio of 1.

### Methylated DNA Immunoprecipitation (MeDIP) and Real-Time qPCR

This method allows studies about DNA modifications and was performed as in [67]. Briefly, 1 μg of denatured and sonicated genomic DNA was immunoprecipitated overnight with 1 μg of rabbit anti 5mC antibody in 100μL of IP buffer (10mM Na-Phosphate pH7, 140mM NaCl, 0,05% TritonX-100). Then, samples were incubated with 10 μL of Dynabeads protein A beads for two hours (Thermo Fisher), washed four times with IP buffer, and bound DNA eluted by incubation with 20 μg proteinase K (Roche) at 50°C in proper digestion buffer. Eluted DNA was recovered and purified using QIAquick PCR purification kit (Qiagen). DNAme on specific TSS were analyzed by qPCR using the following primer pairs:

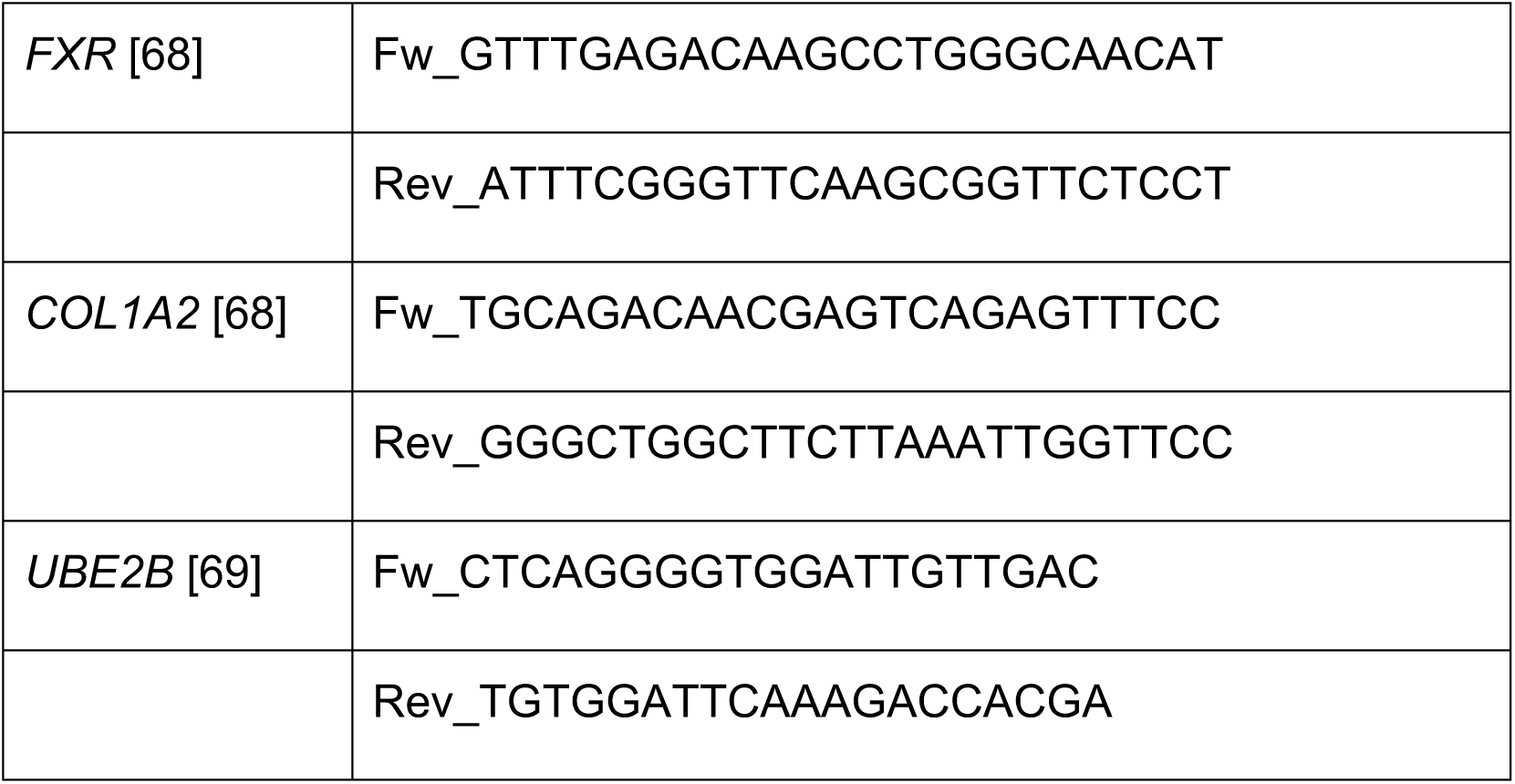

### Dot Blot

5mC dot blot was performed as in [70] with minor modifications. Briefly, genomic DNA was extracted from cells using a NucleoSpin Tissue kit (Macherey-Nagel, 740952), diluted in TE buffer and denatured in 0.4 M NaOH, 10mM EDTA at 95°C for 10 min, then neutralized 1:1 with 2M cold ammonium acetate pH 7 on ice. 500ng DNA were spotted on a PVDF membrane, air dried and baked at 80°C for 10 min, blocked with 5% non-fat milk in TBS-0.1% tween for 1 h, incubated with 5mC primary antibody (1:1000; A-1014 Epigentek) 1h and after three washes in TBS-0.1% tween, incubated with the HRP-conjugated mouse secondary antibody. Signal was acquired on a Chemidoc (BioRad) and band quantified by densitometry with Fiji.

### Combined Bisulfite Restriction Analysis (COBRA)

500 ng of genomic DNA was bisulfite converted with the EpiTect® Fast 48 DNA Bisulfite Kit (Qiagen) according to the manufacturer’s instructions. PCR amplification of converted DNA was performed with platinum Taq DNA polymerase (Thermo Fischer Scientific) and primer pairs adapted from [44] with adjusted annealing temperatures. PCR amplicons were digested with 10 U BstUI (*TDRD6*, Alu), or BstBI (SatII) for 2 h at 65°C and 60°C, respectively. An equal amount of PCR product for the undigested Control was used. Reactions were loaded onto a 3% agarose gel and visualized on a Chemidoc (BioRad). Bands were densitometry quantified by ImageJ and the percentage of methylated DNA was calculated over the total intensity of all the quantified bands.

### 5mC methylation array and analysis

Genomic DNA was extracted from cells using a NucleoSpin Tissue kit (Macherey-Nagel, 740952), DNA concentrations were determined in duplicate using the Quant-IT kit (ThermoFisher). Samples with discordant results were verified in a 2^nd^ series of measurements. DNA quality and correspondence with sample characteristics were evaluated on a subset of the samples by migrating a small amount of DNA on a TapeStation 4200 (Agilent) to calculate the DNA Integrity number (DIN), by a PCR amplification test (simultaneous amplification of 2 microsatellites markers) and a PCR-based verification of the sex of individuals. One microgram of genomic DNA was bisulfite converted with the EpiTect® Fast 96 DNA Bisulfite Kit (Qiagen) according to the manufacturer’s instructions. Genome-wide DNA methylation was analysed on 850000 CpGs with the Infinium Human MethylationEPIC Kit (Illumina) according to the manufacturer’s instructions.

Raw data were extracted using the Bioconductor ChAMP package version 2.18 [71, 72]. All 16 samples were below the quality threshold of less than 10% of failed detection p-value (>0.01) probes (global mean of 0.619% of failed probes) and no sample was therefore removed. Probes were further filtered as follows: detection p-value > 0.01 (23598 probes removed); beadcount <3 in at least 5% of samples (18674 probes); non CpG probes (2761 probes); multi-hit probes as described in [73] (8375 probes).

Beta values, i.e. methylation level using the ratio of intensities between methylated and unmethylated alleles (between 0 and 1 with 0 being unmethylated and 1 fully methylated) were normalized using Subset-quantile within array normalization (SWAN) normalization [74].

Different methylated probes (DMPs) between samples were obtained with Bioconductor ChAMP package function champ.DMP which uses limma package to calculate differential methylation probes between two phenotypes. To define DMP, a delta of 0.3 of beta-values median (interpreted as 30% methylation difference) with a p-value <0.05 was set. P-values were adjusted with Benjamini-Hochberg method. Heat maps were generated on R software. As stated in the figure legends, heatmaps show a number of DMPs resulting from applying different Δβ values in order to accommodate a reasonable number of probes for graphic representation.

The different DMPs list of pairwise comparisons were annotated for gene locations according to Illumina annotation (Promoter, TSS, gene body, 3’UTR, etc). To discover the epigenetic context and the repetitive sequence feature of DMPs from pairwise comparisons, the Chromatin State Segmentation by HMM from ENCODE/Broad (Broad ChromHMM data [75]) and Repeat Masker [46] (http://www.repeatmasker.org), respectively, were exploited. Among the available ChromHMM datasets, the one relative to HepG2, having the most similar profile to DLD-1 cells, was used. Since the original 4^th^ and 5^th^, 6^th^ and 7^th^, 14^th^ and 15^th^ chromatin states are identical (namely, strong enhancers, weak enhancers and Repetitive/CNV, respectively), they have been merged in our analysis for a total of 12 chromatin states. Statistical analysis were performed using Chi square test. Statistically changing categories, i.e. the one contributing the most to the p-val are indicated with “*” and assessed based on the standardized residuals, stdres> 2).

### Hi-C processing and data analysis

Hi-C assays were performed with an Arima Hi-C plus kit (Arima Genomics #A510008) according to manufacturer’s Instructions starting from 2*10^6^ cells/reaction. Libraries preparation were performed with a Kapa Hyper Prep Kits with KAPA Library Amplification Primer Mix (Roche, 07962347001), quantified by Qubit (Invitrogen), checked on a Tapestation device (Agilent) and Pair-End sequenced on a NovaSeq platform (Illumina).

#### Hi-C data processing

Hi-C libraries were processed using the distiller pipeline (https://github. com/open2c/distiller-nf). Briefly, Hi-C sequencing reads were mapped to the hg38 assembly using bwa mem, and alignments were processed, deduplicated and classified into contact pairs using pairtools (https://github.com/open2c/pairtools). Hi-C pairs were aggregated into multiresolution contact matrices, filtered and normalized by iterative correction using the cooler package. Genome browser plots were generated using HiGlass and coolbox. Contact frequency vs distance profiles were calculated using cooltools.

#### IPG analysis

Spectral clustering and analysis were performed on Hi-C from untreated *^NA^DNMT1* cells using the inspectro package as described in [28] applied to the first 10 eigenvectors of the contact matrix at 50kb resolution. We compared the results of k-means with several functional and epigenomic tracks, including H3K9me3 and DNAme (this study), H3K27me3 and RNA-seq [58], as well as GC content and distance from centromere. Based on cluster metrics and interpretability, we selected k=9 clusters and combined two pairs and one triple of clusters whose functional profiles were largely similar but differed in overall proximity to the centromere to yield 5 IPGs. Comparison of IPGs with labels from SNIPER was generated using bioframe. Summaries of IPG-level observed/expected contact frequency were generated using cooltools.

### ChIP-seq and RNA-seq data processing

ChIP-seq data, including data from [58] were processed following the steps of the ENCODE ChIP-seq pipeline (https://github.com/ENCODE-DCC/chip-seq-pipeline2) with slight modifications using a simplified custom snakemake workflow as described in [28]. ChIP-seq tracks were compared by aggregating bigwig signal at 10-kb resolution using pybbi and generating density scatter plots using the matplotlib extension for datashader. RNA-seq data from [58] was processed with the nf-core RNA-seq pipeline using default parameters.

## SUPPLEMENTARY FIGURE LEGENDS

**Supplementary Figure 1.**
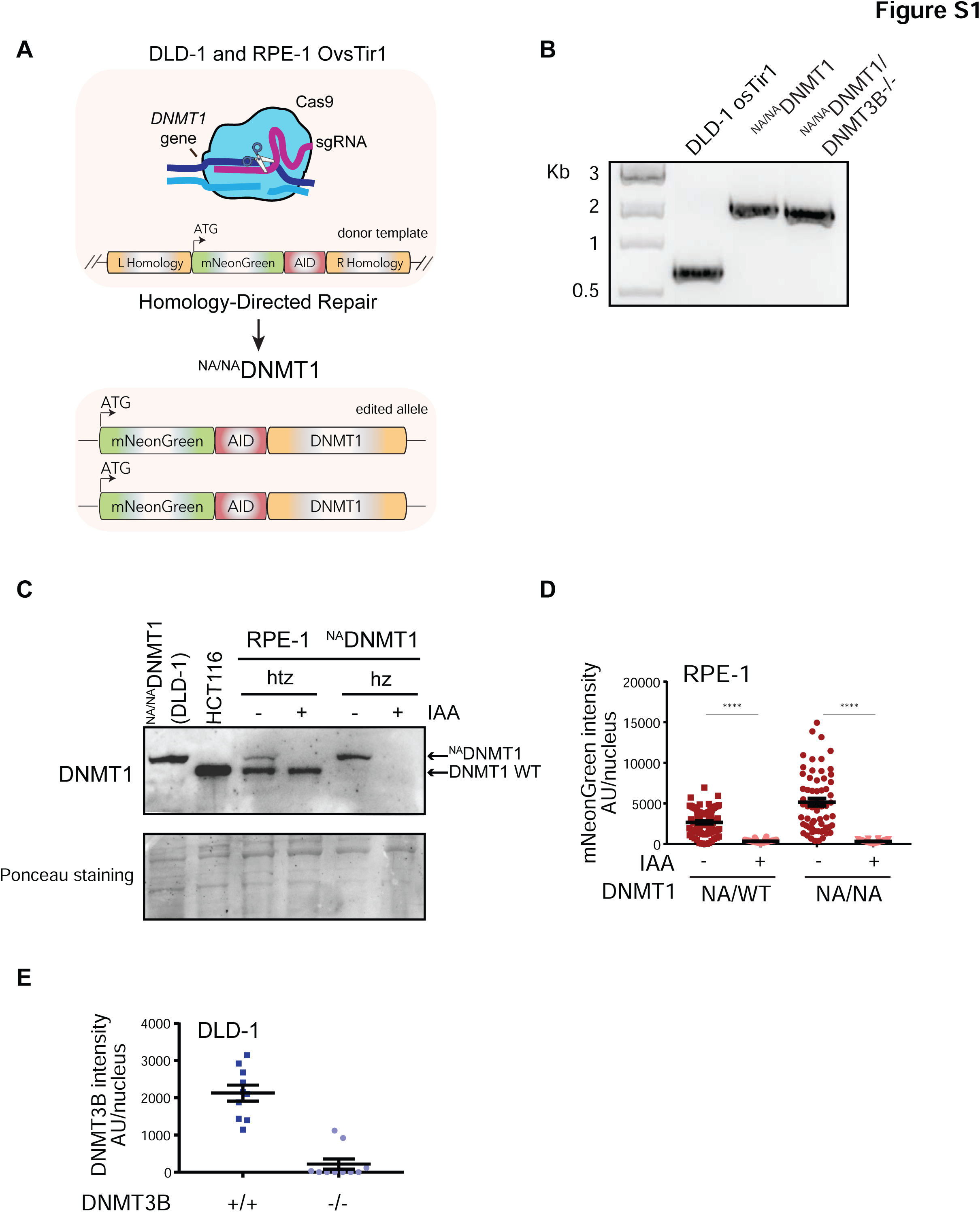
Generation of an inducible DNMT1 degradation system. **A.** Schematics of the endogenous DNMT1 gene tagging strategy by CRISPR/Cas9 approach. **B.** PCR on selected wild type, *^NA^DNMT1* and *^NA^DNMT1/DNMT3B^-/-^*clones showing homozygous tagging of DNMT1 gene with the mNeonGreen-AID module. **C.** Immunoblot analysis with DNMT1 antibody of the indicated cell lines. IAA treatment: 24 h; htz: heterozygous; hz: homozygous. Ponceau staining is shown as loading control. **D.** Quantification of mNeonGreen fluorescence signal in the indicated cell lines. IAA treatment: 24 h. Each dot represents one analyzed nucleus (N > 60 for conditions). Error bars represent the SEM. Unpaired t test: ****p <0.0001. **E.** Quantification of DNMT3B nuclear signal in *DNMT3B* WT and KO cell lines. Each dot corresponds to one analyzed nucleus. Each dot represents one analyzed nucleus (N =10 cells). Error bars represent SEM.

**Supplementary Figure 2.**
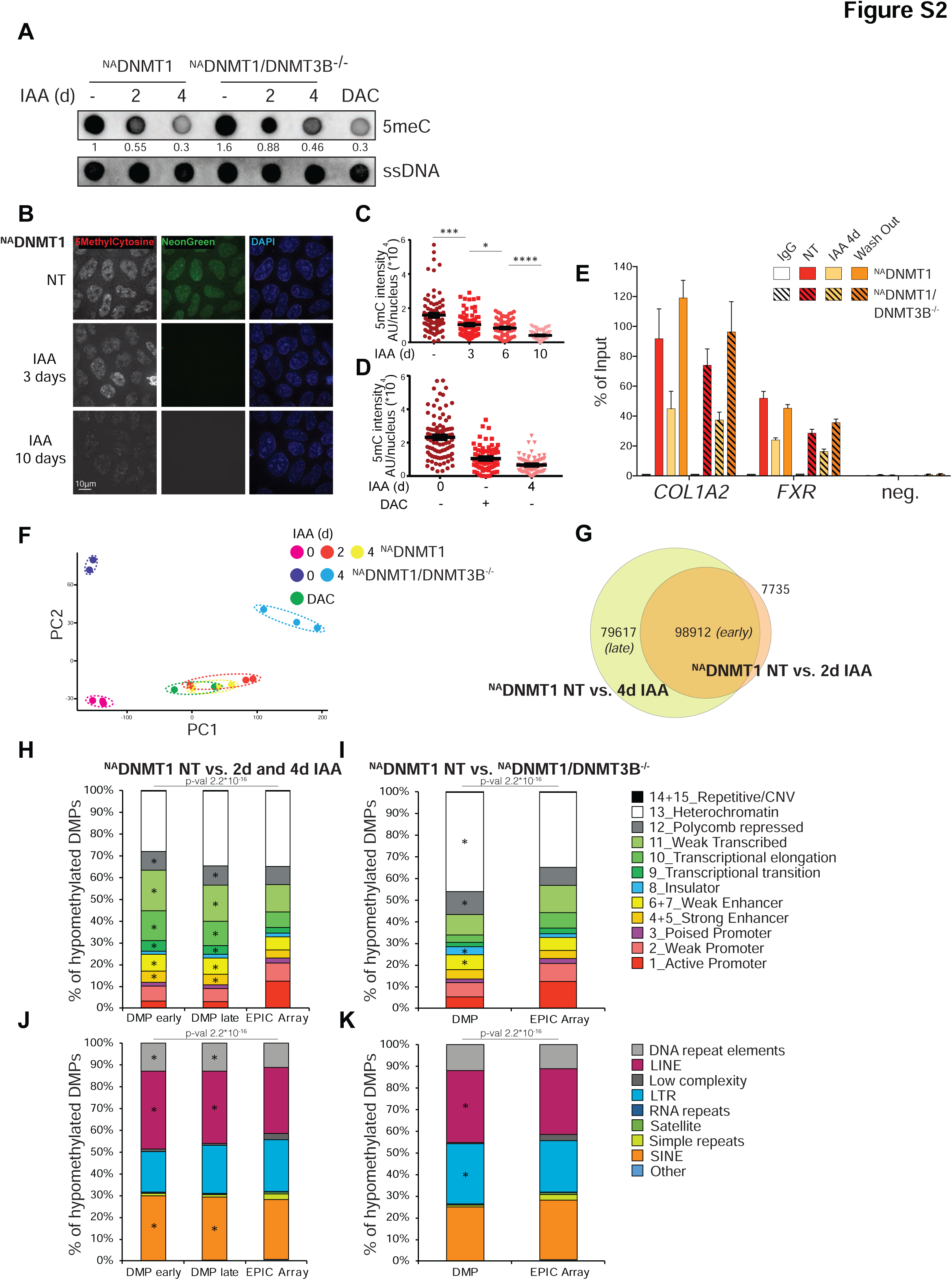
Induced DNMT1 degradation leads to progressive DNA demethylation. **A.** Dot blot analysis with a 5meC antibody of DNA from the indicated cell lines and conditions. Single strand DNA (ssDNA) served as loading control. DAC treatment: 4 days, 2.5µM. Numbers indicates densitometry quantification of signal intensity of 5meC normalized to ssDNA. **B.** Representative immunofluorescence images of the indicated cell lines and treatments with a 5meC antibody. mNeonGreen was used to detect DNMT1. Scale bar, 10µm. **C.** Quantification of 5meC nuclear signal at different times of IAA treatment of *^NA^DNMT1* cells. Each dot corresponds to one analyzed nucleus. Error bars represent SEM. Unpaired t test: * p = 0.0348, *** p = 0.002 ****p <0.0001. **D.** Quantification of 5meC nuclear signal of 4 days IAA-treated *^NA^DNMT1* cells compared to DAC treatment (4 days, 2.5µM). Each dot corresponds to one analyzed nucleus. Error bars represent SEM. **E.** Methylated DNA Immunoprecipitation (MeDIP) analysis at selected promoter regions in the indicated cell lines and conditions. IgG served as isotype control. **F.** Principal Component Analysis (PCA) plot of DNAme variations among the indicated cell lines and treatments. Biological replicates are represented by individual dot, enclosed by dashed circles. **G.** Venn diagram indicating the numbers of the Differentially Methylated Probes (DMPs, Δβ-value ≥ 30%) identified in the indicated pairwise comparisons. **H-I.** Distribution of the indicated pairwise hypo-DMPs relative to the 15 chromatin states (ChromHMM from ENCODE, Hep2G cell line) compared to the distribution of all the probes present on the EPIC Array. Since the described 4^th^ and 5^th^, 6^th^ and 7^th^, 14^th^ and 15^th^ chromatin states (strong enhancers, weak enhancers and Repetitive/CNV, respectively) are identical, they have been merged in our analysis for a total of 12 chromatin states. Chi-square test was used to calculate p-value and define significant changes in the distribution of DMPs of the indicated categories relative to EPIC array composition. Stars (*) indicate the chromatin states with major change (i.e. contributing the most to the p-value based on the standardized residuals, stdres> 2). **J-K.** Distribution of the indicated pairwise hypo-DMPs relative to the annotated DNA repeats (Repeat Masker) compared to the distribution of all the probes present on the EPIC Array. Chi-square test was used to calculate p-value and define significant changes in the distribution of DMPs of the indicated categories relative to EPIC array composition. Stars (*) indicate the categories with major change (i.e. contributing the most to the p-value based on the standardized residuals, stdres> 2).

**Supplementary Figure 3.**
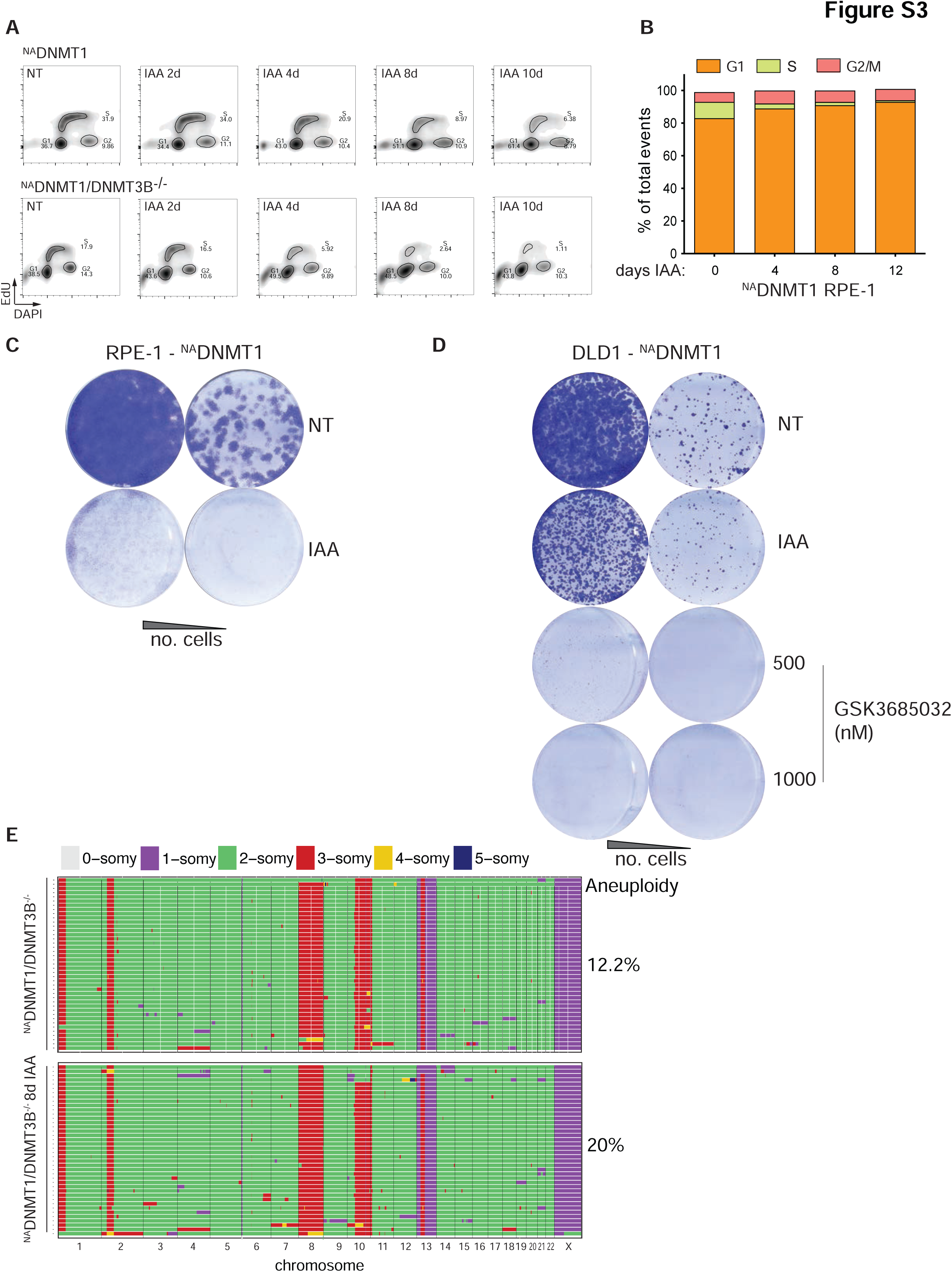
DNMTs degradation impairs cell proliferation. **A.** Representative plot of cell cycle analysis of *^NA^DNMT1* and *^NA^DNMT1/DNMT3B^-/-^*DLD-1 cells upon IAA treatment for the indicated times. Shown gates were set with an auto-gating tool. **B.** Quantification of cell cycle phases of the *^NA^DNMT1* RPE-1 cells treated with IAA for the indicated times. **C.** Representative images of colony formation assay of the *^NA^DNMT1* RPE-1 cells treated with IAA for 12 days. **D.** Representative images of colony formation assay of the *^NA^DNMT1* DLD-1 cells treated with IAA or GSK3685032 for 12 days. **E.** Single cell Whole-Genome Sequencing (scWGS) of *^NA^DNMT1/DNMT3B^-/-^*treated as indicated. Each row represents an individual cell, and chromosomes are plotted as columns. Colors correspond to a defined copy-number state (legend on the right). The percentage of cell showing whole-chromosome aneuploidy (loss or gain, besides chromosome 8 and 13) is also indicated.

**Supplementary Figure 4.**
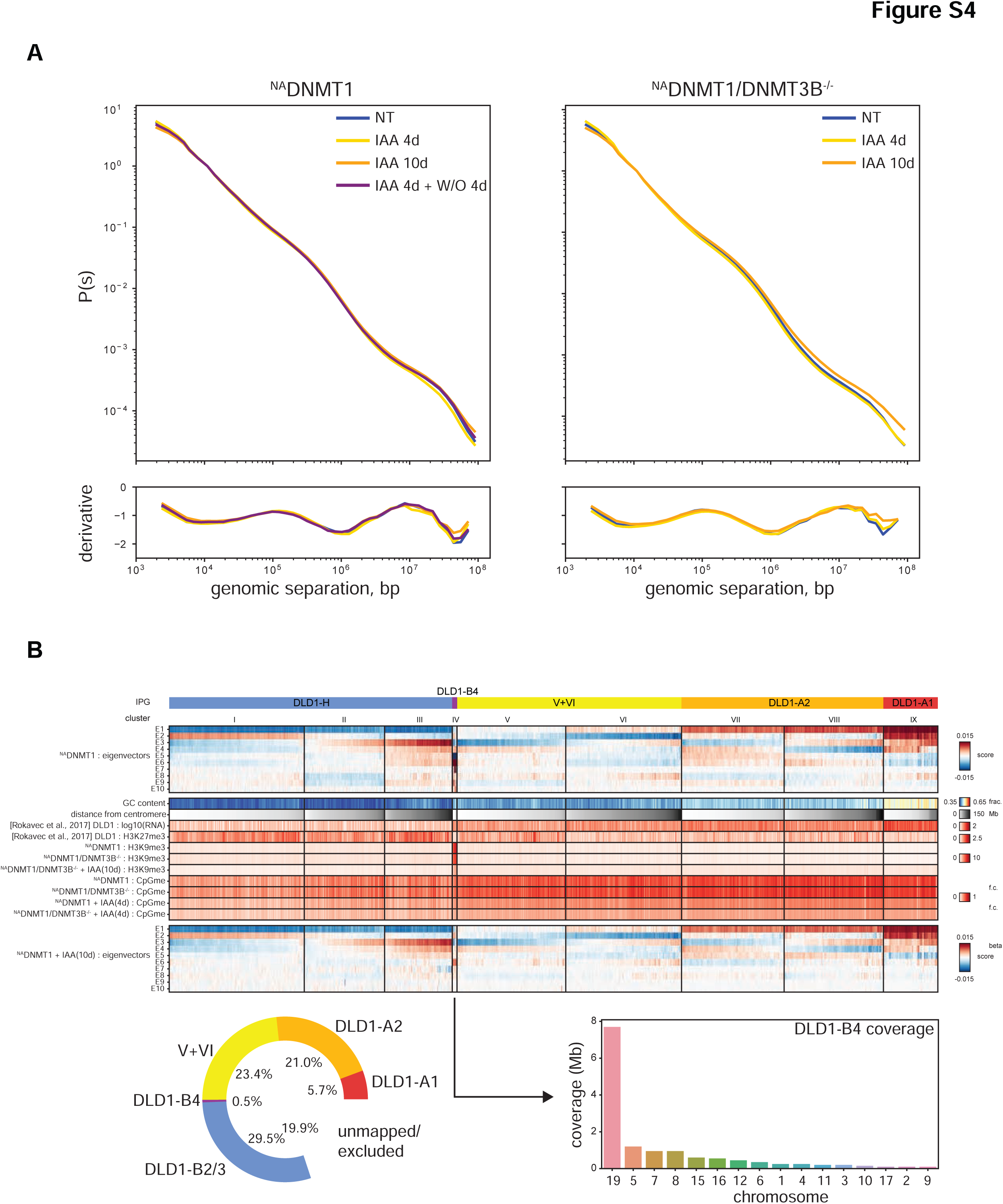
Assignments of subnuclear compartmentalization of *^NA^DNMT1* and *^NA^DNMT1/DNMT3B^-/-^* DLD-1 cells. **A.** Plots showing interaction frequency as a function of genomic distance (P(s) curves) for Hi-C maps performed in the indicated cell lines and treatments. Bottom panel shows the P(s) derivative indicating average size of extruded loops. Washout experiment (W/O) was analyzed 4d after IAA withdrawal (IAA treatment: 4 days). **B.** Integrated heatmaps of various features (rows) for each 50-kb genomic bin (column). Bins are sorted into groups derived from clustering of eigenvectors from untreated *^NA^DNMT1* Hi-C data. Within each cluster, bins are sorted by distance from the centromere. Top section: Consolidated IPG labels (color-coded) and eigenvector clusters labels I-IX below them, followed by the first 10 trans eigenvectors from untreated *^NA^DNMT1* Hi-C. Middle section: Mean signal intensity of functional genomics features: GC content, distance from centromere, bulk RNA-seq signal and H3K27me3 ChIP-seq from wildtype DLD1 cells [58], H3K9me3 ChIP-seq and CpG methylation data from this study. Bottom section: first 10 trans eigenvectors from 10-day IAA treated ^NA^DNMT1 Hi-C data, showing cluster IV (i.e. IPG DLD1-B4) as the most impacted by treatment. Bottom left inset: Donut plot of IPG composition across autosomes, showing 19% of regions excluded from analysis due to poor mappability or translocations. Bottom right inset: Distribution of DLD1-B4 coverage by chromosome shows it is predominantly found on chromosome 19.

**Supplementary Figure 5.**
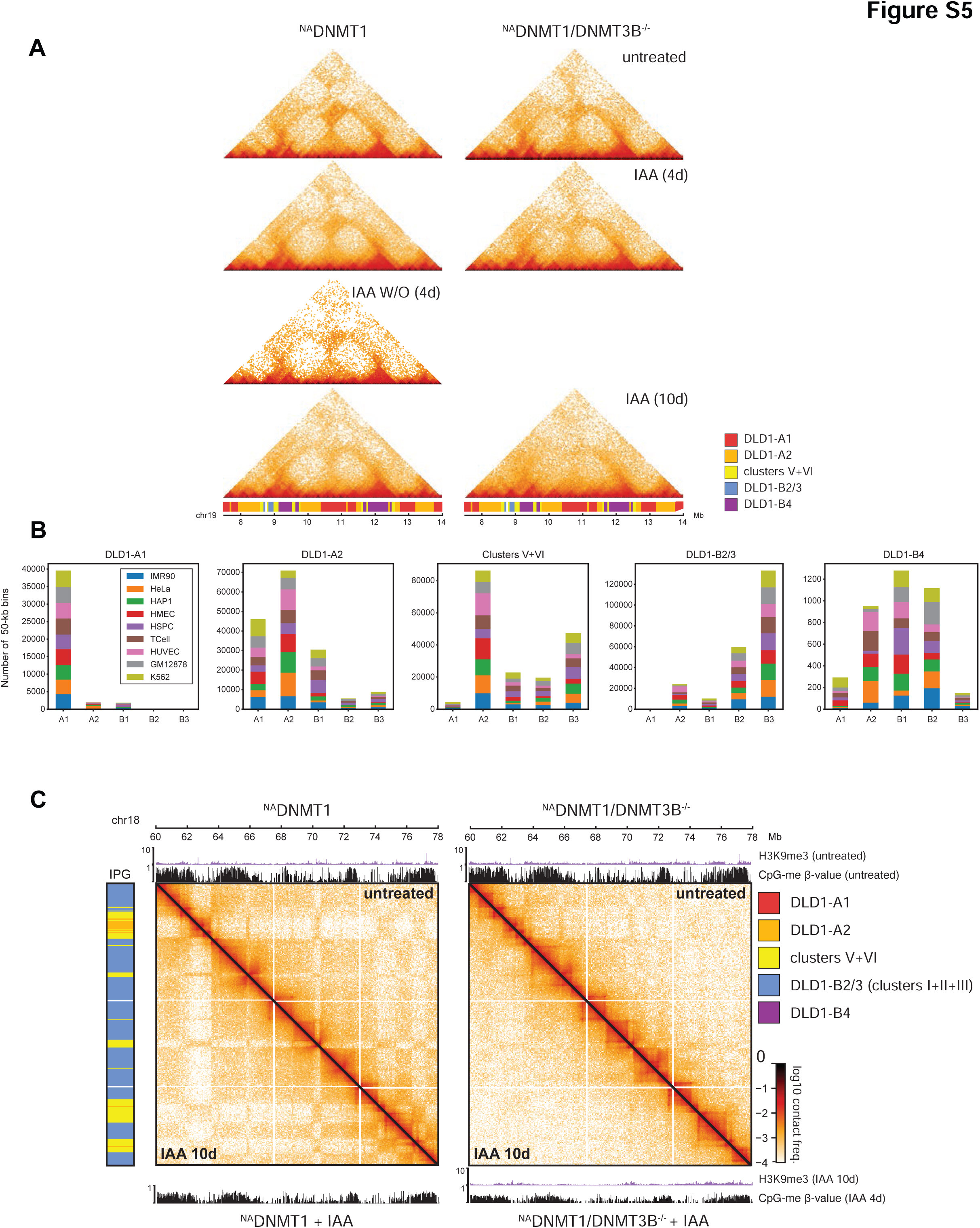
Subnuclear compartmentalization of ^NA^DNMT1 and ^NA^DNMT1/DNMT3B^-/-^ DLD1. **A.** Time courses of Hi-C in the chromosome 19 region from Fig 4E, in the indicated cell lines and conditions. Washout experiment was analyzed 4d after IAA withdrawal (IAA treatment: 4 days). **B.** Distribution of SNIPER subcompartment label predictions of 50-kb loci from untreated *^NA^DNMT1* cells, grouped by the de novo IPG classification in this study. **C.** Example of DLD1-B2/3 interaction profile disruption of DLD1-B2/3 in the *^NA^DNMT1/DNMT3B^-/-^*background compared to ^NA^DNMT1 in a large region on chromosome 18. Relative H3K9me3 and CpG methylation (SWAN normalized β−value from the EPIC array) signals are also shown (as in Fig. 4F).

